# Enhancing Transcriptome Mapping with Rapid PRO-seq Profiling of Nascent RNA

**DOI:** 10.1101/2024.05.08.593182

**Authors:** Pradeep Reddy Cingaram, Felipe Beckedorff, Jingyin Yue, Fan Liu, Helena Gomes Dos Santos, Ramin Shiekhattar

## Abstract

Precision nuclear run-on (PRO) sequencing (PRO-seq) is a powerful technique for mapping polymerase active sites with nucleotide resolution and measuring newly synthesized transcripts at both promoters and enhancer elements. The current PRO-seq protocol is time-intensive, technically challenging, and requires a large amount of starting material. To overcome these limitations, we developed rapid PRO-seq (rPRO-seq) which utilizes pre-adenylated single-stranded DNAs (AppDNA), a dimer blocking oligonucleotide (DBO), on-bead 5’ RNA end repair, and column-based purification. These modifications enabled efficient transcriptome mapping within a single day (∼12 hours) increasing ligation efficiency, abolished adapter dimers, and reduced sample loss and RNA degradation. We demonstrate the reproducibility of rPRO-seq in measuring polymerases at promoters, gene bodies, and enhancers as compared to original PRO-seq protocols. Additionally, rPRO-seq is scalable, allowing for transcriptome mapping with as little as 25,000 cells. We apply rPRO-seq to study the role of Integrator in mouse hematopoietic stem and progenitor cell (mHSPC) homeostasis, identifying *Ints11* as an essential component of transcriptional regulation and RNA processing in mHSPC homeostasis. Overall, rPRO-seq represents a significant advance in the field of nascent transcript analyses and will be a valuable tool for generating patient-specific genome-wide transcription profiles with minimal sample requirements.

## Introduction

In recent years, next-generation sequencing (NGS) technology has revolutionized our understanding of transcription by enabling precise mapping of various RNA species, ranging from messenger RNAs (mRNAs) to long intergenic noncoding RNAs (lincRNAs). Transcriptional regulatory elements, including distal and proximal control regions, regulate the RNA polymerase II (RNAPII) transcription cycle, which includes transcription initiation and RNAPII pause-release^1^. Although RNA sequencing (RNA-seq) has facilitated the discovery of novel transcripts and the identification and annotation of various classes of cellular RNAs, it primarily measures steady-state, mostly cytoplasmic, RNA species. Steady-state RNA levels are a consequence of the equilibrium between the synthesis rate, RNA processing, and RNA degradation, leading to the failure of RNA-seq to detect many low-abundant and unstable RNAs that fall below its detection threshold^2,3^. For example, biologically active enhancer RNAs (eRNAs) and some other long noncoding RNAs (lncRNAs), are not easily detected with these approaches^4,5^. Beyond RNA-seq’s limitations, RNAPII ChIP-seq enables genome-wide identification and quantification of both transcriptionally active and inactive RNAPII. However, this technique lacks strand specificity, making it difficult to determine the transcriptional status and direction of RNAPII. Furthermore, compared to RNA-seq-based methods, ChIP-seq gives relatively high background and heavily relies on antibody specificity, making it challenging to quantitate RNAPII density at genes and their regulatory elements.

To overcome these limitations, several groups have developed methodologies to measure dynamic changes in RNAPII using Nuclear Run-On (NRO)-based methods, including Global run-on sequencing (GRO-seq)^6^ and Precision nuclear run-on (PRO) sequencing^7^. Additionally, complementary methodologies of native elongating transcript sequencing (NET-seq)^8^, and transient transcriptome sequencing (TT-seq)^9^ have been developed. These methods measure transcription by profiling nascent RNAs elongated by polymerases and accurately map the polymerases and their start sites. PRO-seq represents a genome-wide improvement of the NRO assay that enable the identification of both initiating and elongating RNAPII with single nucleotide resolution at high sensitivity and specificity. However, PRO-seq presents technical challenges that require several days of hands-on experimentation, resulting in a significant amount of adapter dimer, and demand a large starting material of 1–2 × 10^7^ cells^10^.

Here we present a rapid Precision Run-On and sequencing (rPRO-seq) technique that produces a robust mapping of the nascent transcriptome with Illumina-compatible sequencing libraries within a single day (∼12 h). We applied rPRO-seq to measure nascent RNA in HeLa cells which provided results with a similar quality as that of the conventional PRO-seq method. The rPRO-seq library process uses pre-adenylated single-stranded DNAs (P-3’ App-DNA) as a substrate in a 3’ adapter ligation reaction to prevent unwanted ligation products. Furthermore, to avoid adapter-dimer formation, we introduce a dimer blocking oligonucleotide (DBO) into the reaction mixture after the 3’ adapter ligation step and before the 5’ adapter ligation step, such that the 3’ adapter is no longer capable of ligating to a 5’ adapter. We also demonstrate that rPRO-seq can be performed with as few as 25,000 cells, potentially extending the studies of nascent RNA in model systems with a limited cell number. In this study, the application of rPRO-seq not only reveals the crucial role of INTS11 in transcriptional regulation and RNA processing in mHSPC, but also showcases the potential of rPRO-seq as a significant advance in analyses of nascent transcription.

### Design

The rapid Precision nuclear Run-On and sequencing (rPRO-seq) protocol represents a significant improvement over conventional PRO-seq methodology, tailored to overcome inherent technical complexities and time constraints. Central to its design are strategic modifications aimed at maximizing library efficiency while minimizing hands-on processing time. A defining feature of rPRO-seq is its innovative integration of pre-adenylated DNA (P-3′ App-DNA) adapters and a dimer blocking oligonucleotide (DBO) within the ligation process. This strategic incorporation circumvents undesirable side product formation, including self-circularization and off-target RNA species ligation. The innovative integration of pre-adenylated DNA (P-3′ App-DNA) adapters and a dimer blocking oligonucleotide (DBO) into the ligation process effectively circumvents undesirable side product formation, such as self-circularization and off-target RNA species ligation. Additionally, the incorporation of on-bead 5’ RNA end repair and subsequent column-based purification streamlines library preparation, allowing for completion within a 12-hour timeframe, compared to several days required for conventional PRO-seq. This streamlined approach offers an expedient and technically tractable method for high-resolution mapping of nascent transcriptomes, enhancing the efficiency and precision of genomic research.

## Results

### Detailing the rPRO-seq protocol

We present rapid Precision nuclear Run-On and sequencing (rPRO-seq) protocol (Figure 1A), that addresses challenges associated with traditional PRO-seq methods, such as formation of adapter-dimers, an extended time required to prepare libraries and the reduced efficiency when using limited input material. The conventional PRO-seq protocol entails several days and involves several phenol:chloroform extractions and ethanol precipitation steps. The new rPRO-seq protocol utilizes pre-adenylated DNA (P-3′ App-DNA) adapters during the ligation process to reduce side product formation, thereby preventing self-circularization, self-polymerization, and ligation to RNA species other than the adapter in the sample. In addition, P-3′ App-DNA adapters are used during the 3′ adapter ligation step to perform the subsequent ligation in the absence of ATP to avoid ATP-dependent side reactions^11,12^ (Figure 1B). This approach enables ligation solely between the 5′ end of the P-3′ App-DNA adapter and the 3′-OH of the sample RNA, thus preventing ATP-dependent circularization and multimerization of RNAs with 5′ phosphorylated ends^13^.

**Figure 1.**
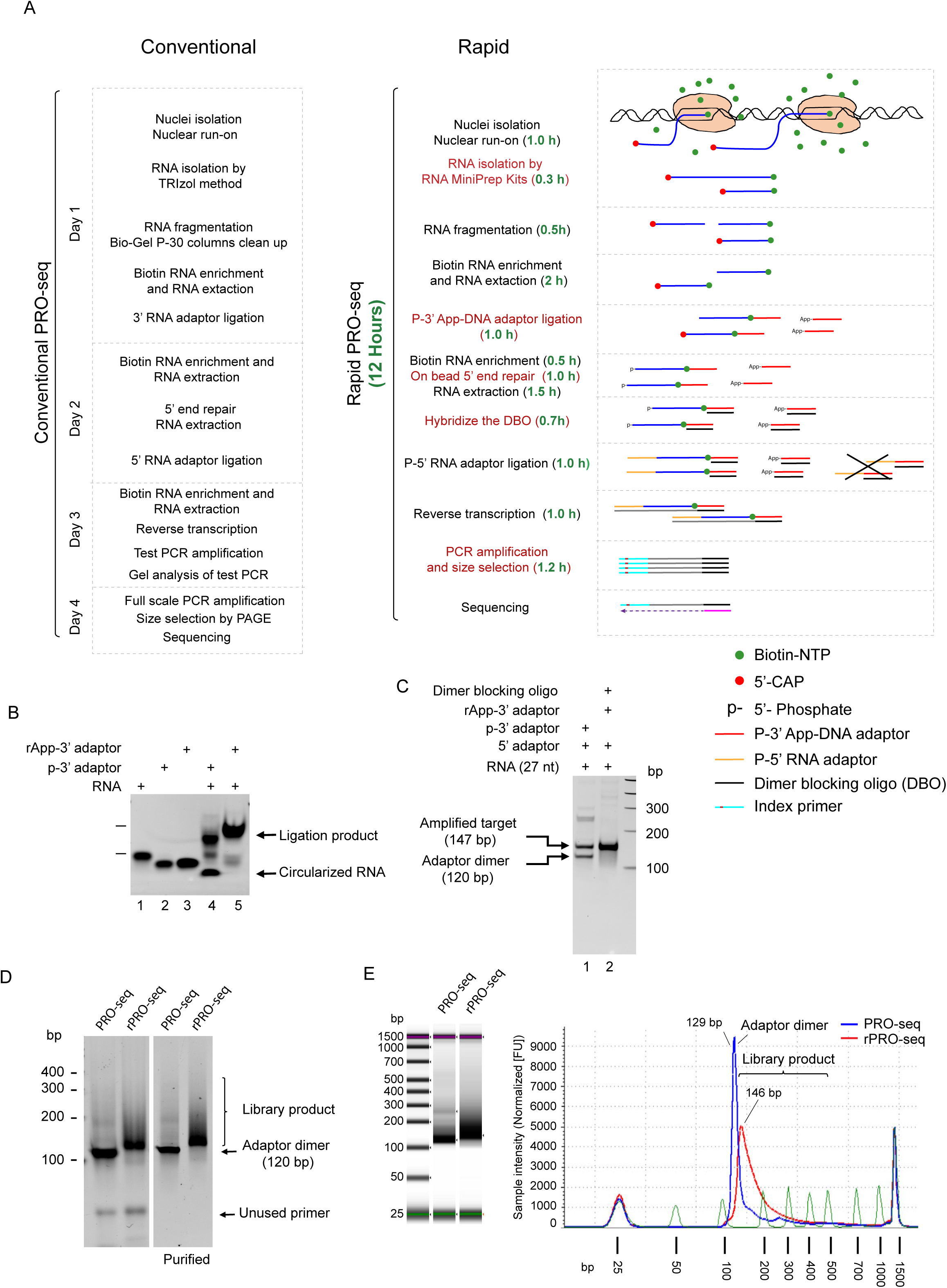
Schematics of the rPRO-seq method. (A) Flowchart of the conventional PRO-seq and rPRO-seq procedures. (B) Shows a 5% native polyacrylamide gel electrophoresis (PAGE), ligation of non-adenylated (p-3’ adapter) or 3’ pre-adenylated (App-3’ adapter) adapter to the synthetic phosphorylated target RNA using T4 RNA ligase (lane 4) or T4 RNA ligase 2 in absence of ATP (lane 5). (C) shows a 5% native PAGE. Lane 1 shows PCR amplified DNA derived from synthetic phosphorylated target RNA after adapter ligation using normal methods and reverse transcription and PCR amplification. An indicated band corresponding to an amplified adapter dimer (120 bp) and band at the position (147 bp) corresponding to an amplified target. Lane 2 shows the amplified products obtained using dimer blocking oligonucleotide, a single band related to the amplified product (147 bp) and no band at the position (120 bp) related to an amplified adapter dimer. (D) PRO-seq and rPRO-seq libraries after final amplification, analyzed by 5% PAGE before and after purification. (E) Agilent 4200 TapeStation system screen tape gel image and electropherogram of a representative PRO-seq and rPRO-seq libraries.

In PRO-seq RNA characterization, adapter-dimer formation during the second ligation process is a common issue due to an excess of 3’- and 5’-adapters in the system (Figure 1C). To counter this, we introduce a dimer blocking oligonucleotide (DBO) that hybridizes with any unbound excess 3’ adapter after the 3’-ligation reaction (Figure 1C, D and E). This hybridization transforms the single-stranded DNA adapter into a double-stranded molecule, preventing its ligation to the 5’ adapters during subsequent ligation.

Compared to traditional PRO-seq methods, rPRO-seq offers several advantages. It eliminates the need for numerous phenol:chloroform extractions and ethanol precipitations through the implementation of downstream on-bead 5’ RNA end repair enzymatic reactions. Column-based purification of RNA after the run-on reaction is provided, which eliminates another organic extraction step. Furthermore, the protocol has been improved to eliminate the need for laborious test amplifications or polyacrylamide gel electrophoresis (PAGE) purification. Overall, rPRO-seq can be completed in as little as 12 hours, compared to the 4-to-5-day duration required for traditional PRO-seq.

Our rPRO-seq protocol provides a more efficient and less time-consuming method for PRO-seq library preparation, particularly when input material is limited. The use of P-3′ App-DNA adapters and a DBO during the ligation steps significantly reduces the formation of side products and adapter dimers, respectively. This protocol also reduces the number of extraction and purification steps required and is less laborious than traditional methods, making it a valuable tool for studying gene expression at a genome-wide level.

### Direct comparison of PRO-seq and rPRO-seq using HeLa cells

The conventional PRO-seq method necessitates a significant amount of starting material, making it unsuitable for use with primary cells or cell lines that cannot be cultured in large quantities. To obtain optimal results with conventional Pro-seq, approximately 1-2 x 10^7^ cells are required. In a comparative analysis, we assessed PRO-seq and rPRO-seq using HeLa cells, a widely utilized cancer cell line. Sequences obtained from 10 million cells using conventional PRO-seq were compared to those obtained from 0.5, 0.1, 0.05, and 0.025 ×10^6^ cells (20- to 400-fold fewer cells than used for PRO-seq) using rPRO-seq. We found that as few as 25,000-50,000 cells yielded results using rPRO-seq comparable to that of PRO-seq performed with 10 million cells. The genome browser tracks of EFNA1 and EIF5, two moderately expressed genes in HeLa cells attest to the robustness of rPRO-seq results when using as little as 25,000 cells (Figure 2A and B).

**Figure 2.**
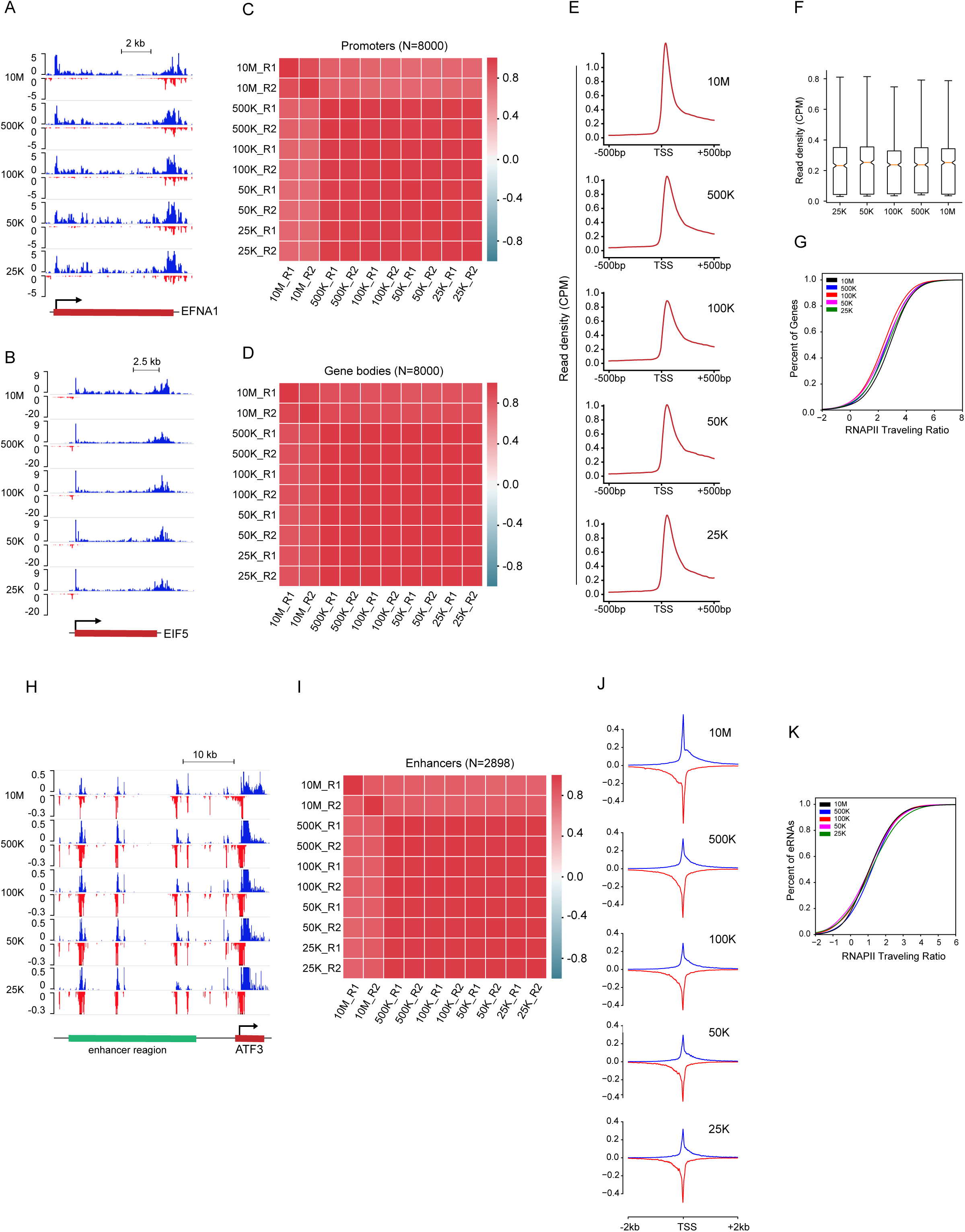
Evaluation between PRO-seq and rPRO-seq in HeLa cells. A and B) Genome browser screenshots of PRO-seq (10M cells) and rPRO-seq (500K, 100K, 50K and 25K cells) at EFNA1 (A) and EIF5 (B). (C) Pearson correlation heatmap of average read density for 8000 active promoters between replicates, protocols and number of cells. (D) Pearson correlation heatmaps of average read density for the gene bodies of 8000 active genes between replicates, protocols and number of cells. (E) Metagene profiles of PRO-seq (10M cells), and rPRO-seq signal at promoters of expressed genes (n = 8000) using different number of cells. TSS, Transcription Start Site. (F) Boxplots of promoter read density at expressed genes in PRO-seq (10M) compared to rPRO-seq (different number of cells). No significant differences in mean read density at promoters compared to the reference 10M (by two-sample t-test). (G) RNAPII raveling ratio (TR) at expressed genes in PRO-seq (10M) and rPRO-seq (different number of cells). (H) Genome browser track example of an enhancer region (eRNA). (I) Pearson correlation heatmaps of the average read density per eRNA (n = 2898) between replicates and protocols. (J) PRO-seq and rPRO-seq profiles of signal at eRNAs using different number of cells. (K) RNAPII raveling ratio at eRNA regions.

We obtained a set of transcriptionally engaged RNAPII regions, including 8,000 transcriptionally active protein coding genes and 2,898 enhancers expressing enhancer RNAs (eRNAs) published using the traditional PRO-seq method on HeLa cells^14^. To assess rPRO-seq’s reproducibility and to compare it with the traditional PRO-seq method, we measure the density of PRO-seq reads at these promoter regions and gene bodies. We then performed Pearson correlation analyses using the average read density per promoter (Figure 2C) and gene body (Figure 2D) between multiple replicates and protocols. We find a high pairwise correlation between multiple replicates using the two protocols (Promoters; r ≥ 0.7389 and Gene bodies; r ≥ 0.8477). When comparing the normalized read density, we find similar profiles at promoters (defined here as −500 nt from TSS to +500 nt) (Figure 2E) and gene bodies (Figure 2F) across protocols. We next determine the RNAPII Traveling Ratio (TR) also referred to as Pausing Index (PI) (the read ratio between the proximal promoter and the region corresponding to the gene body) at 8,000 actively transcribed genes (Figure 2G). Traveling Ratios display no discernible differences between multiple replicates or the two protocols.

We aimed to extend the scope of our findings, moving beyond the exploration of protein coding genes to include enhancer regions. The genome browser tracks, ATF3 enhancer RNAs (eRNAs) present evidence for the robustness of rPRO-seq results, even with a minimal cell count of 25,000 (Figure 2H). In our rigorous examination of rPRO-seq’s reproducibility and its comparison with the traditional PRO-seq method within enhancer regions, we meticulously quantified the density of PRO-seq reads at 2,898 enhancer sites. Advancing further, we applied Pearson correlation analyses using the average read density across multiple replicates and protocols (Figure 2I). The results revealed a strong pairwise correlation (eRNAs; r ≥ 0.7670), indicating consistency across multiple replicates using the two protocols. Upon comparing the read density profiles at enhancers (defined as −2000 nt from the transcription start site (TSS) of enhancers to +2000 nt), we noted consistent patterns across different protocols (Figure 2J). Furthermore, our examination of Traveling Ratios at 2,898 enhancers (Figure K) revealed no notable differences among multiple replicates or between the two protocols. Taken together, analyses of nascent transcription at promoters, gene bodies and enhancers reveal that rPRO-seq provides reproducible results using as little as 25,000 cells, extending nascent transcriptome analysis to model systems where low cell numbers precluded nascent transcription analysis.

### Utilizing rPRO-seq to assess transcriptome changes in mouse hematopoietic stem and progenitor cells (mHSPC)

We next performed rPRO-seq in mHSPC cells to evaluate its sensitivity and ability to measure nascent transcripts using limited cell numbers, given that HSPCs are a rare cell type comprising only 0.01% of nucleated cells in bone marrow^15^. Additionally, we investigated the changes in nascent transcription in HSPCs using a conditional knockout mouse for *Ints11*, the catalytic subunit of Integrator complex. This allowed the determination of Integrator contribution to transcription at promoters and enhancers of mHSPCs. To gain insight into the transcriptional targets regulated by *Ints11*, we isolated cKit+ lineage negative cells from *Ints11^WT/WT^* and *Ints11^Δ/Δ^* mice HSPC populations^16^. Approximately, 500,000 cells were used for these experiments to obtain optimal results since overall level of transcripts in HSPCs were lower than that of HeLa cells. *Ints11* knockout resulted in the differential expression of 3525 genes, and nearly half of them were either down- (1624 genes) or up-regulated (1901 genes) (Figure 3A–C). There was a substantial change in rPRO-seq signals associated with the reads in the body of the genes after integrator knockout (Figure 3D). The analysis of down-regulated transcripts through Gene Ontology revealed a significant impact on signal-responsive pathways such as mRNA processing, IL-6 signaling, EGFR1 signaling, insulin signaling, and NF-kB signaling pathways (Figure 3E).

**Figure 3.**
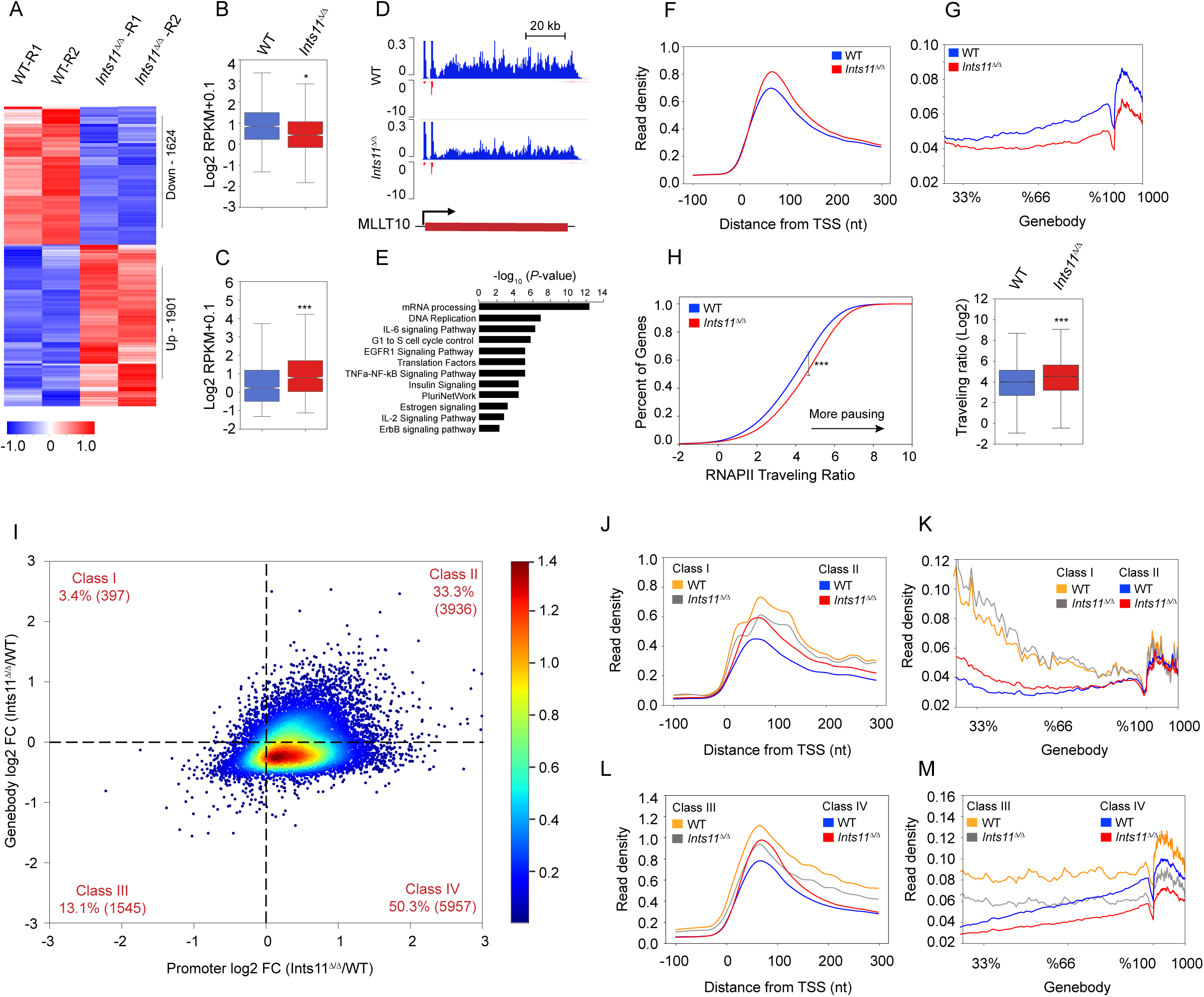
Integrator facilitates transcriptional elongation in mHSPC. (A) Heatmaps of rPRO-seq signal for differentially expressed genes in mHSPC WT and Ints11^Δ/Δ^. (B) and (C) Boxplot analysis of rPRO-seq coverage (RPKM) in gene bodies in down regulated (n=1901) (B) and up regulated (n=1624) (C) genes. t-test, *p < 0.05, ***p < 0.0001. (D) Representative tracks of rPRO-seq signal in mHSPC cells at MLLT10. (E) Gene Ontology analysis of the 1624 transcripts downregulated by Ints11-depletion, as defined from rPRO-seq data. (F) and (G) rPRO-seq profiles at the promoter (F) and gene bodies (G). (H) rPRO-seq traveling ratio at all expressed genes in mHSPC WT and Ints11^Δ/Δ^. Average profile KS test, box plot t-test, ***p < 0.0001. (I) Two-dimensional traveling matrix (TM) comparing fold change of rPRO-seq signals at promoters (x axis) and gene bodies (y axis) by comparing Ints11^Δ/Δ^ with WT. (J) and (K) Metagene profiles of rPRO-seq promoter read density and gene body profile at class I and II genes in mHSPC WT and Ints11^Δ/Δ^. (L) and (M) Metagene profiles of rPRO-seq promoter read density and gene body profile at class III and IV genes in mHSPC WT and Ints11^Δ/Δ^.

To investigate the role of Integrator complex more thoroughly in transcriptional dynamics, we divided the rPRO-seq signal into reads covering the promoter region which corresponds to transcriptional initiation and those at gene body regions corresponding to transcriptional elongation. Consistent with our previous results following knock-down of INTS11 in human cancer cell lines^17–19^, depletion of INTS11 in HSPCs, leads to increased RNAPII read density at the promoter region and reduced read density through the gene body, resulting in decreased productive transcriptional elongation (Figure 3F and G). We next determine traveling ratio (TR) for RNAPII defined as rPRO-seq read densities of the promoter regions divided by reads at the gene bodies. We find the TR profile moving to the right in *Ints11^Δ/Δ^* HSPCs, suggesting increased pausing in the absence of Integrator with a concomitant decreased productive transcriptional elongation (Figure 3H).

To simultaneously interrogate the alterations at promoters and gene bodies of individual INTS11-responsive genes, we used the traveling matrix (TM) which was previously developed to determine the two-dimensional Gaussian distribution of RNAPII^17^ (Figure 3I). TM resolves changes in promoters and gene bodies of individual genes and these changes are then represented as two-dimensional density plot. While deletion of INTS11 results in widespread transcriptional effects, in concordance with the TR profile, 50% of genes on the TM display an increased reads at their promoters concomitant with decreased read density in their gene bodies reflective of increased pausing and diminished transcriptional elongation (Figure 3I).

To analyze the behavior of genes on the traveling matrix in more detail, the genes on the matrix were separated into four gene classes (class I–IV), representing differentially expressed genes with concurrent promoter and gene body signal alterations (Figure 3I-M). We assessed RNAPII promoter and gene body profiles to further validate how INTS11 regulates transcription of the four gene classes determined by the TM (Figures 3J-3M). INTS11depletion altered the coordinates of many genes in the TM, but most INTS11-responsive genes appeared in class II and IV classes, reflecting increased RNAPII pausing (Extended Figure 3A-D). Furthermore, the evident enrichment of genes in class IV, where increased promoter signal is accompanied by reduced reads at gene bodies, suggests that INTS11 is required for RNAPII pause-release and productive elongation at these loci. Complementing the traveling matrix, analyses of TR profiles revealed a pronounced shift to the right for class IV genes reflecting a prominent elongation defect (Extended Figure 3D). Taken together, INTS11 loss elicits RNAPII pausing with reduced elongation, as shown by class IV genes in TM.

### Assessing eRNAs using rPRO-seq

Beyond measuring protein coding gene expression, PRO-seq is an important tool to assess enhancer RNA expression. We next used rPRO-seq reads at intergenic sites that display histone H3 lysine 27 acetylation to determine eRNAs at active enhancers using established pipelines (Fig.4A). Since Integrator plays a role in eRNA processing^17,20^, we also sought to determine alternations in eRNAs expression after deletion of *Ints11* using rPRO-Seq in mHSPC cells. Using rPRO-seq reads, we determined 3293 enhancer RNA sites in mHSPCs following the pipeline in figure 4A, excluding regions overlapping with annotated transcription start sites (TSS) (Figure 4A). Two examples of active enhancers expressing eRNAs are shown in figure 4B and 4C.

**Figure 4.**
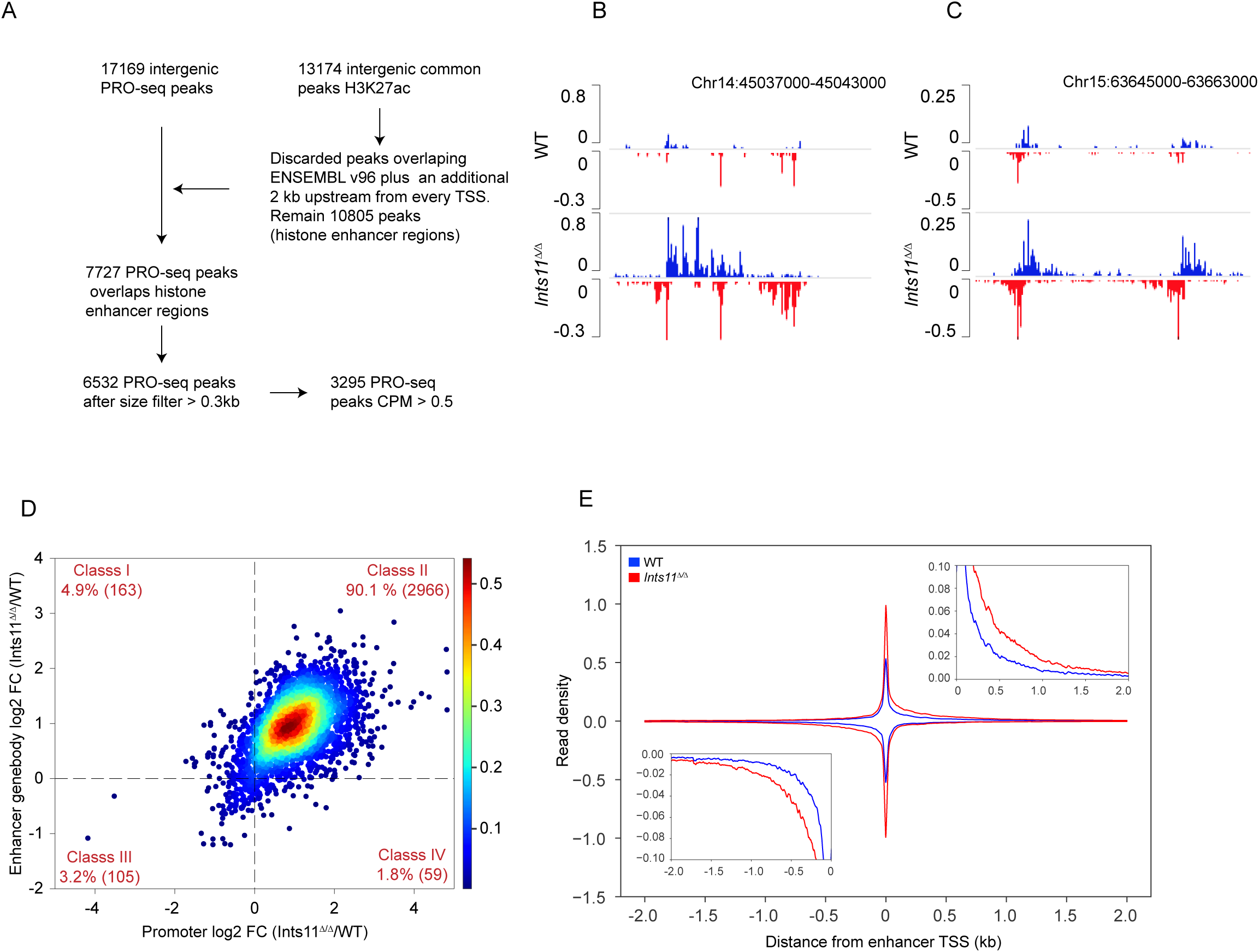
Enhancer transcription regulation by Integrator in mHSPC. (A)Schematic representation of the enhancer analysis for obtaining the set of actively transcribed eRNAs in mHSPC cells. (B) and (C) Genome browser snapshot examples of rPRO-seq signal at enhancers in mHSPC cells. (D) Enhancer Two-dimensional traveling matrix (TM) of rPRO-seq signal in WT and Ints11^Δ/Δ^. (E) rPRO-seq enhancer profile (n = 3293) in mHSPC WT and Ints11^Δ/Δ^.

The rPRO-seq TM and average profiles at enhancers demonstrated augmented transcription, which was explained by increased RNAPII density at the TSS and in the body of eRNA transcripts (Figure 4D and E). Notably, the differential TM profiles at enhancers displayed a markedly distinct pattern compared to the differential TM profiles observed at promoters (Figure 4D). In fact, nearly 90 percent of eRNAs revealed a substantial increase in RNAPII levels near eRNA transcription sites promoters following *Ints11* deletion, accompanied by elevated RNAPII levels within the gene bodies of these eRNAs (class II transcripts, Figure 4D). Specifically, INTS11 depletion resulted in extended signal corresponding to impaired termination at enhancers (Figure 4E). These analyses clearly demonstrated that the impact of *Ints11* loss on RNAPII transcribing eRNAs differed from its effects on mRNAs (class II vs. class IV accumulation, respectively). Overall, these results provide strong evidence that Integrator plays a crucial role in the regulation of eRNA transcription, acting as a promoter-proximal termination complex, which restricts the release of paused RNAPII into the transcript body.

## Discussion

Nascent RNA sequencing is a widely used approach to investigate regulatory steps of RNA polymerase II, such as pausing, elongation, and termination, as well as to obtain accurate quantitative data on gene expression^6–9^. In recent years, researchers have focused on understanding the role of unstable and low-abundant RNA categories in epigenetics and transcription regulation. For instance, low-expressed and poorly conserved long intergenic non-coding RNAs (lincRNAs) have been shown to regulate a variety of biological processes by associating with either repressing or activating chromatin complexes. NRO-based nascent RNA sequencing is a powerful strategy that can map and measure gene expression and the activities of transcriptional regulatory elements, such as enhancers simultaneously.

The current PRO-seq methods have limitations that include technical challenges, a requirement for a large amount of starting material, and the generation of significant amounts of adapter dimers, making it a time-consuming and labor-intensive process. To address these limitations, we have developed a rapid PRO-seq (rPRO-seq) protocol based on nascent RNA-sequencing assays. This new method offers a fast, sensitive, and simple approach to generate genomic libraries of nascent RNA. To avoid unwanted ligation products, we utilized AppDNA as substrates in a 3′ adapter ligation reaction during the preparation of rPRO-seq libraries. Furthermore, to eliminate adapter-dimer formation, a dimer blocking oligonucleotide was added to the reaction mixture after 3’ adapter ligation and before 5’ adapter ligation, thereby preventing the ligation of 3’ adapters to 5’ adapters. Our rPRO-seq method allows for comprehensive and seamless analysis of RNAPII activity in both abundant RNA species such as protein-coding genes, as well as difficult-to-detect RNAs including eRNAs, lincRNAs, and antisense promoter transcripts. Traditional PRO-seq procedures require 4-5 days to process samples, whereas our rPRO-seq protocol offers a significant reduction in processing time to only 12 hours. This is achieved by using AppDNA, dimer blocking oligonucleotides, on-bead 5’ RNA end repair enzymes, and column-based RNA purification. Our rPRO-seq method is a promising approach for investigating nascent RNA transcription in various biological systems.

Our study investigated the performance of a rPRO-seq protocol in comparison to the conventional PRO-seq method. We observed no major differences in promoter or enhancer profiles between the two protocols, as indicated by the aggregate profiles shown in Figure 2. Even with 25,000 cells was low, we still observed transcription at many enhancers. Furthermore, we analyzed the RNAPII Traveling Ratio (TR), also known as Pausing Index (PI), at protein-coding genes (8,000) or enhancers (2,898). Our results show that there were no major differences in promoter, gene body or enhancer profiles across protocols, suggesting that rPRO-seq can reliably quantify promoter, gene body, or enhancer RNA polymerases. Additionally, we demonstrated that rPRO-seq can be performed with less input material than traditional nascent RNA sequencing methods, which typically require between ten and twenty million cells per sample. We performed a serial downscale of rPRO-seq using a modified protocol that utilized AppDNA and a dimer blocking oligonucleotide. Our findings indicate that rPRO-seq can be performed with as few as 25,000 cells, which is 400 times fewer than the input required for PRO-seq. We also successfully applied rPRO-seq to mHSPC, suggesting the broad applicability of this method across different cell types. These results highlight the potential of rPRO-seq as a valuable tool for investigating nascent RNA transcription in diverse biological systems.

Previous studies have demonstrated that depletion of the Integrator catalytic subunit *Ints11* leads to changes in steady-state gene expression^16^. Our investigation identifies *Ints11* as a crucial component of transcriptional regulation and RNA processing in HSC homeostasis. To explore the Integrator’s role in transcription at promoters and enhancers in HSPCs, we utilized a conditional knockout mouse model (*Ints11^Flox/Flox^*) and depleted INTS11, followed by rPRO-seq analysis. We employed a two-dimensional TM to assess positional changes in RNAPII at promoters and gene bodies in response to deletion of *Ints11*. Our findings revealed a significant reduction in RNAPII signal in the gene bodies of nearly two-thirds of differentially regulated genes, indicating that Integrator facilitates transcriptional elongation in mouse hematopoietic cells. We also observed that RNAPII recruitment was deficient in class I and class III genes, while RNAPII was increased in class II and class IV genes. Furthermore, elongating RNAPII was reduced in these classes, as evidenced by lower steady-state levels of RNA. Our investigation also included an analysis of the enhancer TM, which revealed that a substantial contribution by Integrator to transcriptional termination at these regulatory elements. Specifically, our data showed that INTS11plays an important role in promoting elongation at mouse promoters through the dynamic turnover of paused RNAPII. Thus, Integrator appears to regulate transcription at distinct stages of transcription at mouse enhancers and promoters. Overall, our results provide new insights into the regulatory mechanisms of transcriptional elongation and RNA processing in hematopoietic stem cells.

In summary, the rPRO-seq method offers a fast, sensitive, and simple approach to obtain accurate quantitative data on gene regulation, including unstable and low-abundant RNA categories. This standardized and scalable technique can be applied to a wide range of experimental settings to study the role of transcriptional regulation in various biological processes.

## Limitations

While rPRO-seq has shown success in detecting mRNA and eRNA transcription expression using libraries from a small number of cells, such as 25,000 in cancer cell lines, one should be mindful that studying RNAPII regulatory steps (such as pausing, elongation, and termination) with fewer than 50,000 cells may not provide a very detailed picture of the transcriptional cycle. This mainly results from a limited genomic coverage and library complexity which is alleviated by using a higher cell number. Even at high sequencing depths, rPRO-seq’s single base resolution on a genome-wide scale may lead to relatively low read counts on individual pausing sites, limiting the sensitivity and precision with 25,000 cells. However, this limitation may be overcome by considering and analyzing ensembles of pausing sites.

Additionally, similar to conventional PRO-seq, rPRO-seq selectively captures actively elongating RNA polymerase molecules, potentially missing pre-initiation complexes and some stalled polymerases. The short run-on length used for high-resolution mapping in rPRO-seq may result in the omission of RNA polymerases near the transcription start site (TSS). RNA polymerases positioned very close to the TSS cannot be fully detected as very short reads are removed before alignment, owing to the insufficient length of nascent RNA for unique genome mapping. Furthermore, while rPRO-seq demonstrates reproducibility and yields comparable results to the traditional PRO-seq, its specificity and accuracy may vary depending on the experimental conditions and the biological system under investigation mainly due to the overall strength of transcriptional outputs in individual cell types.

## Acknowledgements

We thank the Shiekhattar lab for constructive discussions and suggestions for experimental design. We thank the Oncogenomics and flow cytometry shared resources at Sylvester Comprehensive Cancer Center for performing high-throughput sequencing. This work was supported by funding from Sylvester Comprehensive Cancer Center P30CA240139 and grants R01 GM078455, and DP1 CA228041 from the National Institute of Health to R.S. Research reported in this publication was supported by the National Cancer Institute of the National Institutes of Health under Award Number P30CA240139. The content is solely the responsibility of the authors and does not necessarily represent the official views of the National Institutes of Health.

## Author Contributions

R.S. and P.R.C. conceived and designed the overall project. P.R.C. performed experiments. J.Y. and F.Liu performed mHSPC isolation experiments. F.B., H.G.D.S. and P.R.C. performed bioinformatics analyses. R.S. and P.R.C. wrote the manuscript with critical feedback from all co-authors.

## EXPERIMENTAL MODEL AND SUBJECT DETAILS

### Cell Lines

HeLa cells were cultured in Dulbecco’s Modified Eagle’s Medium (DMEM) supplemented with 10% FBS at 37°C with 5% CO2.

### Method Details

#### Reagents

TRIzol Reagent (Ambion, Cat no. 15596018), TRIzol LS Reagent (Ambion, Cat no. 10296028), Biotin-11-ATP (PerkinElmer, cat. no. NEL544001EA), Biotin-11-CTP (PerkinElmer, cat. no. NEL542001EA), Biotin-11-GTP (PerkinElmer, cat. no. NEL545001EA), Biotin-11-UTP (PerkinElmer, cat. no. NEL543001EA), Streptavidin M280 beads (Invitrogen, cat. no. 112.06D), SUPERase-In RNase Inhibitor (20 U/μL) (Catalog number: AM2696), T4 RNA Ligase 1, High Concentration (NEB, cat. no. M0437M). RNA 5’ Pyrophosphohydrolase (RppH) (NEB, cat. no. M0356S) can be used with ThermoPol Reaction buffer (NEB, cat. no. B9004S). T4 polynucleotide kinase, (PNK) (NEB, cat. no. M0201L), Superscript III reverse transcriptase (Invitrogen, cat. no. 56575), Deoxynucleotide (dNTP) Solution Mix (NEB, cat. no. N0447S), NEBNext Ultra II Q5 Master Mix (NEB, cat. no. M0544S).

#### Isolation of Mouse Hematopoietic Stem and Progenitor Cells (mHSPC)

To investigate hematopoietic stem and progenitor cells (HSPC), we employed an inducible whole-body knockout model. This involved crossing our animal model with Rosa26-CreERT2 (Cre recombinase-estrogen receptor T2) mice obtained from Jackson Laboratory (Stock No: 008463). Both control and Ints11 floxed mice received intraperitoneal injections of tamoxifen (75mg/Kg per day) for five consecutive days to induce the deletion of Ints11. To isolate HSPC cells, we utilized the Direct Lineage Cell Depletion Kit and cKit (CD117) MicroBeads from Miltenyi Biotec, following the manufacturer’s instructions. In brief, BM cells were initially stained with a cocktail of lineage marker antibodies directly coupled to MicroBeads and then passed through a column. Subsequently, the cells were further purified using cKit MicroBeads to enrich hematopoietic stem and progenitor cell populations.

#### ChIP-Seq data analysis

ChIP-seq datasets from Choukrallah et al.^21^, (GSE60005) and Raw FASTQ data were processed with Trimmomatic v0.32 to remove low-quality reads and then aligned to the human genome hg19 using STAR aligner v2.5.3a^22^. Peaks were called using MACS2.1.2^23^ with the following parameters --shift 75 --nomodel --extsize 200 for narrow peaks. Peaks with fold change > 3 and a q-value < 0.01 were used for downstream analysis. Homer annotate Peaks^24^ v4.9.1-5 was used for peak annotation.

#### PRO-seq, rPRO-seq and data analysis

We conducted PRO-seq according to previously described^10^. For rPRO-seq experiments were conducted following established protocols (see detailed protocol in supplementary file 1). Initially, a mixture of HeLa or mHSPC nuclei and Drosophila S2 cell nuclei as spike-ins was prepared. Nuclear run-on assays were executed by incubating nuclei at 30°C for 3 minutes with 25 μM Biotin-11-ATP/UTP/CTP/GTP (PerkinElmer). Subsequently, total RNA was extracted using Direct-zol RNA Miniprep and fragmented with 0.2 M NaOH for 10 minutes on ice, followed by purification of biotinylated nascent RNAs using streptavidin beads M-280 (Invitrogen). The 3’ RNA adaptor ligation process employs pre-adenylated single-stranded DNAs (P-3’ App-DNA), and T4 RNA ligase 2, truncated KQ, is utilized, incubated at 25°C for 1 hour. Additionally, on-bead 5’ RNA end repair involved using RppH (NEB) to remove the 5’ cap and repair the triphosphate, followed by PNK-mediated repair of the 5’ hydroxyl. The hybridization of DBO involves incorporating the dimer blocking oligonucleotide (DBO) into the RNA within the DBO mix. This is followed by a thermocycler program and subsequent combination with the 5’ adaptor ligation mixes in the RNA DBO mix, with a 1-hour incubation at 25°C. After adapter ligation, SuperScript III reverse transcription was employed. PCR amplification generated 140–350 bp libraries, which were size-selected using AMpure XP beads and sequenced using single-read runs on a NovaSeq 6000.

Raw fastq data were processed as described previosly ^17^. Briefly PRO-seq reads were trimmed by Cutadapt 1.14^25^ and Trimmomatic v0.32^26^ and aligned to the human genome (hg19), or mouse genome (mm10) or the drosophila genome (dm3) by bowtie 1.1.2^27^ PRO-seq signal was normalized by the number of reads mapped to spike-in dm3 genome, and then converted to bigwig data, which were used for downstream analyses. In HeLa cells (human), we used the previously published and extensively studied set of actively transcribed promoters of protein-coding genes and enhancer RNAs^17^. To determine the accurate TSS for each gene in mouse, we separated the mapped reads from PRO-seq by strand and individually call peaks using MACS2.1.2^23^ with the following parameters: --nomodel, --extsize 200 and –narrow to identify narrow peaks. We merged the common peaks from each sample (WT and Ints11 KO) for each stranded peak and used BedTools v2.28^28^ command closest to annotate the peaks against mouse Gencode (v25). Peaks with distance >100nt from an annotated TSS region were discarded to avoid multiple assignments to the same promoter. When multiple TSS from the same reference were detected in the same PRO-seq peak, we selected the most upstream isoform for analysis. Also, when a gene has multiple isoforms, the distance between these TSS was > 350nt and they were assigned to different PRO-seq peaks, we kept all isoforms for analysis. Next, we calculated the RPKM values in promoter region (-50 nt from TSS to + 300 nt) and the region corresponding to the gene body (+301 nt from TSS to TES) for all transcripts used in the mouse analysis (n = 11,835). These values were used as input for the traveling matrix (TM). The TM can be visualized as a 2D distribution by using the log2 ratio (RPKM Ints11 KO over RPKM Ints11 WT) of the promoter region (X axis) and gene body (Y axis). The relative difference generated by this 2D representation can be categorized in four different groups: Class 1: log2 promoter > 0 and log2 gene body < 0; Class 2: log2 promoter < 0 and log2 gene body < 0; Class 3: log2 promoter > 0 and log2 gene body > 0 and Class 4: log2 promoter < 0 and log2 gene body > 0.0 The read counts in the gene body region were used to calculate and determine the differential gene expression (DEG) between conditions using DESeq2^29^. Differentially expressed genes were identified by using a 1.25-fold change cutoff (FDR<0.05) and applying a minimum gene expression filter for the final set of differentially expressed protein-coding genes (0.3 RPKM in both replicates in at least one of the conditions). Enrichment analysis was performed on the differential expressed genes by Enrichr^30,31^.

PRO-seq signal was used to calculate the traveling ratio (TR) as previously described^32^. Briefly, we calculated the ratio between the RPKM values in promoter region (-50 nt from TSS to + 300 nt) over the region corresponding to the gene body (+301 nt from TSS to TES) for all transcripts used in the analysis.

#### Correlation analysis

The correlation analysis was performed using deeptools 3.1.2^33^. Read counts were extracted from the different bam files by multiBamSummary using the ‘BED-file’ option for the promoters, gene bodies, or enhancer regions. Pearson correlation matrices were obtained by plotCorrelation with the options ‘—skipZeros’ and ‘--removeOutliers’.

#### Metagene profiles

Metaprofiles of stranded data were generated with an in-house script based on pyBigWig included in deeptools 3.1.2^33^ by extracting the normalized reads per position at the regions of interest (promoters or enhancers) from the bigwig files and averaging the signal across replicates using a 1 nucleotide bin size.

| Sample | Total Reads | Reads Mapping to hg19 |
| --- | --- | --- |
| Hela_10M_R1 | 113342117 | 67942325 |
| Hela_10M_R2 | 113609471 | 64157575 |
| Hela_25K_R1 | 37272858 | 28421888 |
| Hela_25K_R2 | 36764231 | 28067114 |
| Hela_50K_R1 | 36802002 | 27788209 |
| Hela_50K_R2 | 31225172 | 23284851 |
| Hela_100K_R1 | 36138804 | 27718026 |
| Hela_100K_R2 | 35794986 | 27431818 |
| Hela_500K_R1 | 35695610 | 27636182 |
| Hela_500K_R2 | 37536713 | 29146089 |

| Sample | Total Reads | Reads Mapping to mm10 |
| --- | --- | --- |
| mINTS11_WT_HSPC_R1 | 133573238 | 76689393 |
| mINTS11_WT_HSPC_R2 | 153564659 | 91783806 |
| mINTS11_KO_HSPC_R1 | 137228688 | 82568174 |
| mINTS11_KO_HSPC_R2 | 171501794 | 97013168 |

## Data Availability

The genome-wide data of this study have been deposited in the NCBI Gene Expression Omnibus (GEO) database with the accession number GSE264740.

## Quantification and Statistical Analysis

Significance was determined by either Student’s t test, non-parametric Wilcox test or Kolmogorov–Smirnov test, as indicated.

---

**Extended Figure 3:**
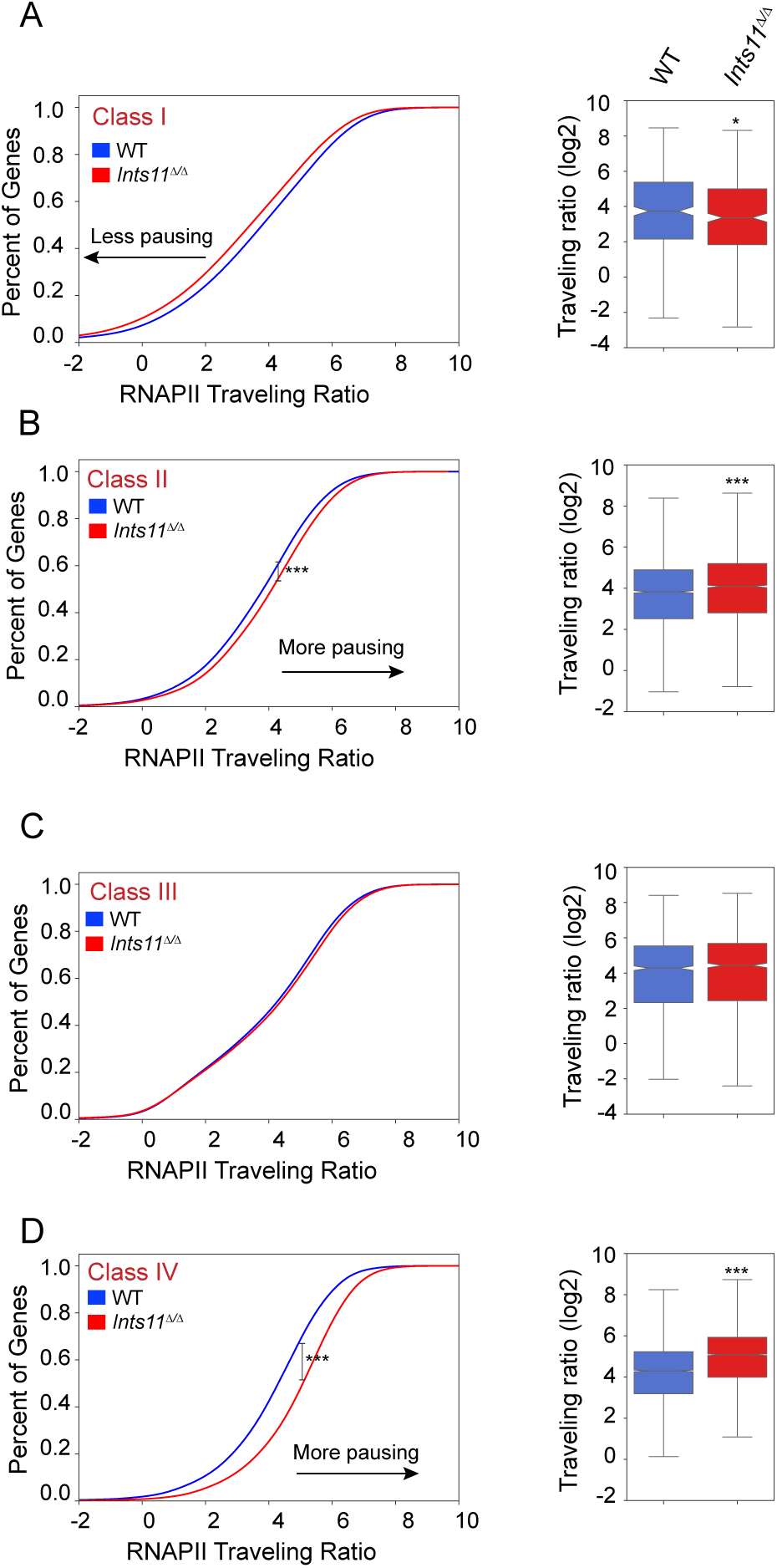
Integrator-mediated enhancement of transcriptional elongation in mHSPC. rPRO-seq traveling ratios of class I genes (A), Class II genes (B), Class III genes (C) and Class IV genes (D) in mHSPC WT and *Ints11^Δ/Δ^*. Average profile KS test ***p < 0.0001, box plot t test *p < 0.05, ***p < 0.0001.

